# Small differences in learning speed for different food qualities can drive efficient collective foraging in ant colonies

**DOI:** 10.1101/274209

**Authors:** F. B. Oberhauser, A. Koch, T. J. Czaczkes

## Abstract

Social insects frequently make important collective decisions, such as selecting the best food sources. Many collective decisions are achieved via communication, for example by differential recruitment depending on resource quality. However, even species without recruitment can respond to a changing environment on collective level by tracking food source quality.

We hypothesised that an apparent collective decision to focus on the highest quality food source can be explained by differential learning of food qualities. Ants may learn the location of higher quality food faster, with most ants finally congregating at the best food source.

To test the effect of reward quality and motivation on learning in *Lasius niger*, we trained individual ants to find a reward of various sucrose molarities on one arm of a T-maze in spring and in autumn after one or four days of starvation.

As hypothesised, ants learned fastest in spring and lowest in autumn, with reduced starvation leading to slower learning. Surprisingly, the effect of food quality and motivation on the learning speed of individuals which persisted in visiting the feeders was small. However, persistence rates varied dramatically: All ants in spring made all (6) return visits to all food qualities, in contrast to 33% of ants in autumn under low starvation.

Fitting the empirical findings into an agent-based model revealed that even a tendency of ants to memorise routes to high quality food sources faster can result in ecologically sensible colony-level behaviour. Low motivation colonies are also choosier, due to increasing sensitivity to food quality.

**Significance statement:** Collective decisions of insects are often achieved via communication and/or other interactions between individuals. However, animals can also make collective decisions in the absence of communication.

We show that foraging motivation and food quality can affect both route memory and the likelihood to return to the food source and thus mediate selective food exploitation. An agent-based model, implemented with our empirical findings, demonstrates that, at the collective level, even small differences in learning lead to ecologically sensible behaviour: mildly starved colonies are selective for high quality food while highly starved colonies exploit all food sources equally.

We therefore suggest that non-interactive factors such as individual learning and the foraging motivation of a colony can mediate or even drive group level behaviour. Instead of accounting collective behaviour exclusively to social interactions, possible contributing individual processes should also be considered.

## Introduction

The ability to choose which resource to exploit is of key importance to both individual animals and animal groups. Groups need to decide where to go while maintaining cohesion, or when to fission or fuse in response to resource availability (Couzin and Krause 2003; Sueur et al. 2011). Eusocial insects are of particular interest for collective decision-making, as they form colonies consisting of many individuals acting as one reproductive unit, which favours the development of elaborate communication systems and food sharing while restricting intra-group conflicts (Ratnieks and Reeve 1992).

Social insects frequently use recruitment to collectively decide on and select a nest site (Mallon et al. 2001; Seeley and Buhrman 2001), to make efficient path choices (Beckers et al. 1992; Reid et al. 2011), and to rapidly exploit food sources (Deneubourg et al. 1983; Seeley et al. 1991; Beckers et al. 1993; Czaczkes and Ratnieks 2012). Recruitment relies on the transmission of social information from one individual to conspecifics, which is often achieved using stereotyped behaviour such as waggle dances in bees (Dyer 2002) or tandem running in ants (Franks and Richardson 2006).

Trail pheromones constitute another way of transmitting location information, and are widely used in ants (Czaczkes et al. 2015b). By depositing pheromone on the substrate on the way back to the nest, successful foragers can both attract their conspecifics and provide an orientation cue to direct them to the food (Wilson 1962; Beckers et al. 1993; Czaczkes and Ratnieks 2012). Naïve ants relying on social information can thus locate new food sources quickly while avoiding trial and error learning and costly mistakes (Galef and Giraldeau 2001) and building up a route memory (Collett et al. 2003). Once a memory is in place, however, it tends to be followed in preference to social information (Leuthold et al. 1976; Aron et al. 1988; Fourcassie and Beugnon 1988; Harrison et al. 1989; Grüter et al. 2008, 2011; Stroeymeyt et al. 2011; Almeida et al. 2017).

Yet even species with no recruitment or communication manage to make sensible decisions at group level and to adjust their behaviour to a changing environment: Cockroaches (*Blattella germanica*) were found to aggregate at food sources based on a retention effect - individuals stayed longer at food sources with high neighbour density (Lihoreau et al. 2010). A similar mechanism was found in greenhead ants (*Rhytidoponera metallica*), a species which does not recruit to food sources. The ants successfully tracked the highest food quality in the environment without comparison between the available food sources. This ability is thought to be achieved by both the tendency of individuals to feed for longer at higher quality food sources as well as a retentive effect of conspecifics to newcomers (Dussutour and Nicolis 2013). As a result, a gradual improvement is observed: While animals are spread to all food sources initially, they concentrate on certain food sources over time until most animals feed on the best food source. This stepwise optimisation process is similar to an annealing process, and represents an efficient and simple solution to optimisation problems (Kirkpatrick et al. 1983; Černý 1985).

Another possible mechanism for driving apparent collective decisions without active recruitment could be based on learning. Many social insects learn the location of food sources very quickly, often after only a single visit (Aron et al. 1988; Grüter et al. 2011) and can have retention times up to months (Salo and Rosengren 2001). They can form associations of odour and food locations and use them to recall and navigate to at least two different feeder locations (Reinhard et al. 2006; Czaczkes et al. 2014). Thus, factors affecting memory formation could have significant impact on the foraging efficiency of a colony. Two particularly ecologically relevant factors are (i) food quality (i.e. reward magnitude) and (ii) motivation (for example starvation level or season), both of which are known to affect collective foraging (Seeley 1986).

An effect of reward magnitude on memory formation is a key part of theories of learning (Rescorla and Wagner 1972), and has been reliably demonstrated in many animals including honey bees. Higher sucrose reward concentrations, for example, increase the probability and retention time of proboscis extension response (PER) conditioning (e.g. Scheiner et al. 1999, 2004, 2005). Honey bees make more return visits to feeders offering sweeter rewards (Seeley 1986) and display higher persistence to return to depleted once-profitable foraging locations which used to offer high concentrations of sucrose (Al Toufailia et al. 2013). Higher persistence leads to more visits to the same food location which facilitates learning and positively influences memory retention (Menzel 1999).

Food deprivation level is known to heavily affect foraging motivation in social insects (e.g. Cosens and Toussaint 1986; Mailleux et al. 2006). Honey bees differ in the amount of sucrose concentration needed to recruit others and have higher thresholds when plenty of food is available (Lindauer 1948; Seeley 1986). Moreover, high satiation levels can disrupt memory formation in honey bees (Friedrich et al. 2004).

Foraging motivation and sucrose thresholds also vary between seasons (Quinet et al. 1997; Ray and Ferneyhough 1997; Beekman and Ratnieks 2000; Scheiner et al. 2003). In temperate regions many animals, including ants (Quinet et al. 1997; Cook et al. 2011), show reduced foraging towards autumn as their reproductive period is over, resources decline (Mailleux et al. 2006) and they prepare to overwinter.

Taken together, we hypothesised that both the motivation of foraging ants, influenced by season and food deprivation levels, as well as the quality of the food source they find, will affect the ant’s route memory formation. High reward and/or high starvation could facilitate rapid learning, while ants at low motivation levels or ants finding low quality food may form memories less rapidly, or be more likely to deviate from their memories. If learning is more likely for higher quality food sources, we hypothesise that ants will tend to memorize and return to higher quality food sources. This should result in an annealing process taking place (Kirkpatrick et al. 1983; Černý 1985): at the beginning, ants forage at all food sources, but as ants finding low quality food are more likely to deviate from their memories, they might end up at the best food source as time progresses. Eventually, most ants should be foraging at the highest quality food source.

Here, we investigated the effect of reward quality and motivation level on memory formation in *Lasius niger* foragers. Reward magnitude was varied by offering different concentrations of sucrose solutions. Motivation levels were varied by testing in spring after four days of food deprivation (high motivation), in autumn after four days of food deprivation (moderate motivation) and in autumn after one day of food deprivation (low motivation). We then used this data to build an agent based model to understand how reward magnitude and motivation level affect the ability of ant colonies to selectively choose the best of multiple foraging locations.

## Material and methods

### (a) Collection and rearing of colonies

Stock colonies of the black garden ant, *Lasius niger*, were collected on the campus of the University of Regensburg, and kept in plastic foraging boxes with a layer of plaster of Paris on the bottom. Each box contained a circular plaster nest (14 cm diameter, 2 cm high). The collected colonies were queenless, and consisted of 500+ workers. Queenless colonies still forage and lay pheromone trails, and are frequently used in foraging experiments (Devigne and Detrain 2002; Dussutour et al. 2004). All colonies were kept in a 12:12 day/night cycle and were provided *ad libitum* with water and Bhatkar diet, a mixture of egg, agar, honey and vitamins (Bhatkar and Whitcomb 1970). The colonies were deprived of food for either 1 or 4 days prior to each trial.

Data was collected in spring (April – May 2016) and in autumn (September – November 2016) to test for seasonal effects (referred to as spring or autumn, respectively). Eight colonies were tested in each experiment. Six of the colonies tested in spring could be retested in autumn. Two of the original colonies were not available for testing, and were replaced by new colonies.

### (b)Experimental procedure

Several ants were allowed onto a plastic T-maze covered with paper overlays via a drawbridge. The runway towards the maze head was tapered to prevent guidance by pheromone before the ant had to choose a side (Fig. 1) (Popp et al. 2017). A sucrose solution droplet (Merck KGaA, Darmstadt, Germany) was presented on a plastic feeder at one side of the T-maze head while a droplet of water was presented on a feeder on the opposite side. The side containing the sugar reward remained the same throughout each test. The T-maze head was recorded from above with a Panasonic DMC-FZ1000 camera, facilitating the observation of ants’ decisions.

**Fig. 1.**
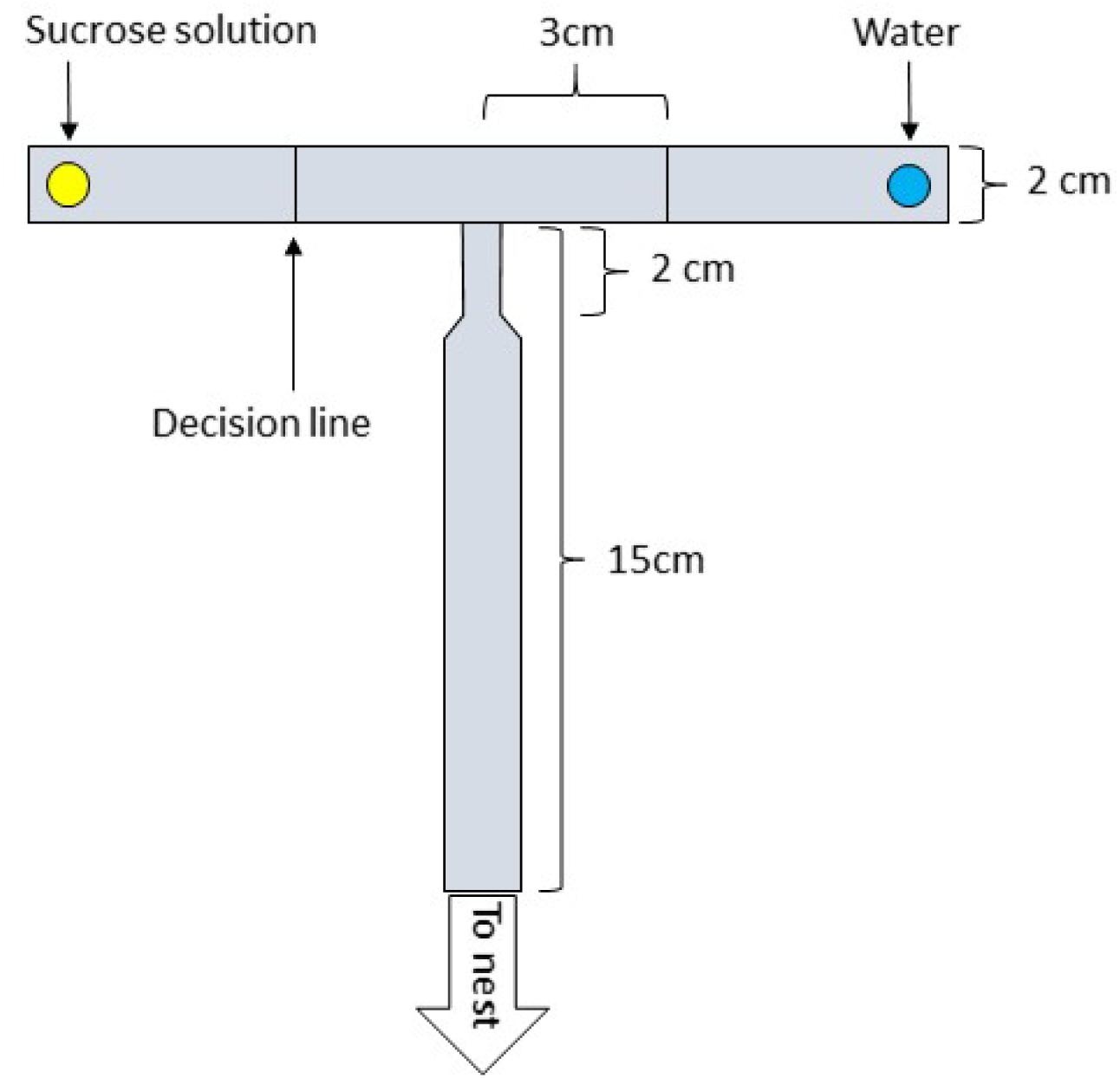
Experimental setup. The ants entered the plastic runway via a moveable drawbridge. The end of the 20cm long runway leading to the maze head was tapered. On one maze arm, a sucrose droplet of either 0.125, 0.5 or 1.5M was presented while a water droplet was presented on the other arm

Each colony was tested with three different concentrations of sucrose solution (0.125, 0.5 or 1.5M). Furthermore, to control for possible side preference, as is often reported in ants (Hunt et al. 2014), each colony was tested on both sides of the T-maze.

The first 6 ants (5 in the autumn low motivation treatment) to reach the feeder were individually marked with a dot of acrylic paint on the abdomen and all other ants were returned to the colony. The drawbridge was used to selectively allow only the marked ants onto the runway thereafter. Each time an ant walked over the maze head the paper overlay was replaced to remove any trail pheromone that might be deposited. On each outward visit for each ant, the first decision line (situated 3cm inwards on each arm, Fig. 1) crossed by the ant with its antennae was scored as its’ initial decision. The first droplet contact on each visit was scored as its’ final decision. The experiments were not conducted blind to treatment, but ants rarely changed arms after their initial decision (see OSM 3 table S3-1), and decisions were unambiguous, thus restricting observer bias. Neither the position of the sucrose solution nor its concentration were changed during each trial. Colonies were tested once per week except for the autumn low starvation experiment, where they were tested once per 2 weeks. All marked ants were permanently removed from the colony after testing to prevent pseudo-replication.

### c)Statistical Analysis

Data were analysed using generalized linear mixed-effect models (GLMM) (Bolker et al. 2009) in R version 3.4.1 (R Core Team 2016). GLMMs were fitted using the lmer function (Bates et al. 2015). As the data were binomial (correct / incorrect), a binomial distribution and logit link were used. Since multiple ants were tested per colony and each ant made repeated visits, we included ant ID nested in colony as random factors. Each model was tested for fit, dispersion and zero inflation using the DHARMa package (Hartig 2016).

The model predictors and interactions were defined *a priori*, as suggested by Forstmeier and Schielzeth (2011), as:

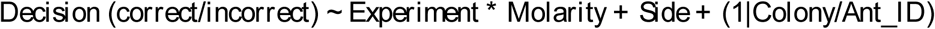

Molarity of the food was included as continuous variable. All p-values presented were corrected for multiple testing using the Benjamini–Hochberg method (Benjamini and Hochberg 1995). In total, 688 ants were tested, 104 of which performed fewer than 6 return visits and were excluded from the analysis to decrease variance in motivation effects, resulting in 584 ants used for analysis. A complete annotated script and output for all data handling and statistical analysis is presented in online supplementary material (OSM) 3. The complete raw data is presented in OSM4.

In only a small fraction (2.09%) of visits, the initial and final decision differed. In most such cases, ants chose the correct side in their initial decision and then U-turned and chose the wrong side on their final decision (table S3-1). For simplicity, due to these small differences, we used only the final decision of each ant as measure of performance in the final analysis.

### d)agent-based model

We developed an agent based model in order to study the effect of individual learning rates on colony-level foraging behaviour. The model is an adaptation of an earlier model of *L. niger* foraging (Czaczkes et al. 2015a) and was coded in Netlogo 6 (Wilensky 1999). Here we provide an overview of the model, a detailed description following the ODD (Overview, Design concepts, Details) protocol (Grimm et al. 2006, 2010) is provided in OSM1, and the annotated model itself is provided in OSM2.

In the model, an ant colony in the centre of the model environment was surrounded by three food sources, either in random positions or in fixed positions equidistant from the nest. The resources were of quality 0, 1, and 2, which respectively represent molarities of 0.125, 0.5 and 1.5. Ants explored the environment using a correlated random walk. If they found a food source, the ants learned the location of the food source and returned to the nest. On returning to the nest, the probability of the ant following its memory or beginning scouting again depended on the motivation level of the colony (a global variable) and the quality of the resource. The probability of memory following was taken directly from the empirical data gathered in our experiments. Memory was modelled in one of two ways. First, we modelled memory in the standard manner, with each ant showing the same behaviour, based on the average behaviour of ants taken from the empirical data (“average ant model”). However, the average behaviour of individuals is a poor description of each individual’s behaviour (Pamir et al. 2011). Thus, we also modelled the ant’s memory use to be variable between individuals, with the average centred on the empirical data, but individuals varying around that (“variable ants model”). This implementation is based on the extended learning curves model of Pamir et al. (2011). The model was run for 3000 time steps, and the proportion of food returned from each food source was recorded at the end of the model run.

## Results

Our statistical model revealed that ants significantly improved their performance (correct decisions) with increasing visits (z = 15.33, p < 0.0001), demonstrating that the ants learned the route to the food source.

Interestingly, reward magnitude, i.e. the molarity of the food, did not significantly affect performance positively (z = 1.88, p = 0.35), although a visual inspection of the data suggests that higher sucrose concentrations led to higher proportions of correct decisions (Fig. 2). Altogether, molarity effects were surprisingly modest, especially at high motivation levels, with 75% accuracy on the first visit even for the lowest molarity at the highest motivation level (see Fig. 2a and Fig. S1).

**Fig. 2.**
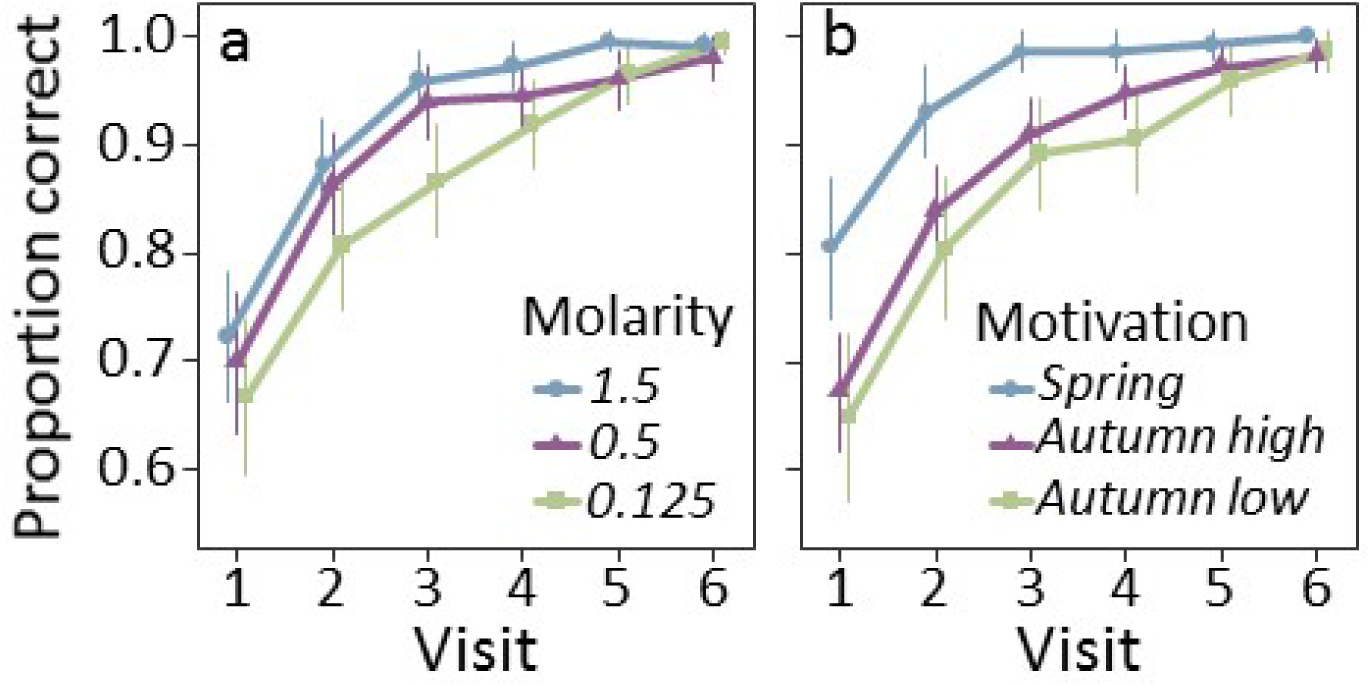
The proportion of correct decisions made by ants as a function of (a) food quality or (b) motivation level. The colonies of the “autumn low” experiment were starved for only 1 day, as opposed to 4 days in the other two experiments. Visit 0 (not shown) is the discovery visit, where ants find the food for the first time. Points show means and error bars show 95% confidence intervals.

It is important to note that we only included ants which finished all 6 visits in our analysis. However, the number of ants which dropped out were not randomly distributed: When presented with 0.125M sucrose, most (>65%) of the ants in the low starvation treatment did not finish all 6 visits (table 1). The number of drop-outs decreased with increasing molarity (∼17% for 1.5M) but remained high in comparison to highly starved autumn or spring ants (<10%, 0%, respectively). By contrast, all ants completed all six visits in spring, irrespective of the molarity, highlighting the importance of motivation on persistence.

**Table 1.** Total number of ants tested and number of ants which finished all 6 visits. Ants with less visits were excluded from the analysis. All ants finished in spring, while many in autumn after 1 day of starvation (autumn low) did not finish.

| Motivation | Molarity | # ants tested | # ants with 6 visits | % dropped |
| --- | --- | --- | --- | --- |
| **Autumn low** | 0.125 | 76 | 25 | **67.1** |
|  | 0.5 | 81 | 57 | **29.6** |
|  | 1.5 | 80 | 66 | **17.5** |
| **Autumn high** | 0.125 | 104 | 98 | **5.8** |
|  | 0.5 | 100 | 91 | **9** |
|  | 1.5 | 103 | 103 | **0** |
| **Spring** | 0.125 | 48 | 48 | **0** |
|  | 0.5 | 48 | 48 | **0** |
|  | 1.5 | 48 | 48 | **0** |
| **Total** |  | **688** | **584** | **15.1** |

Seasonal effects were more prominent: Highly starved ants tested in spring showed faster learning than highly starved ants tested in autumn (z = 4.29, p = 0.0001), and ants tested in autumn that were only starved for one day (z = 5.29, p < 0.0001). No significant difference was found between ants tested in autumn with 4 days of starvation and those starved for one day only (z = 1.88, p = 0.35), although the performance of the 1-day-starved ants tended to be lower (Fig. 2b.).

No significant interactions between motivation (season and starvation level) and reward magnitude (molarity) were found (spring: z = 0.36, p = 0.99; autumn high: z = 0.36, p = 0.99, autumn low: z = 1.67, p = 0.44).

We also noted a side bias, as ants had higher proportions of correct decisions when they had to learn to go to the left (z = 5.59, p < 0.0001). Detailed statistical outputs for all tests are provided in OSM3.

An examination of the individual-level learning rates revealed that most (>70%) ants learned rapidly, already choosing the correct side on the first return visit to the food source (see figure 3A, B). An interesting pattern can be seen when considering the number of successive correct visits: The majority (>75%) of ants in spring made no errors over all 6 visits, while this was only achieved by <60% of ants of the autumn colonies (Fig. 3c, d).

**Fig. 3.**
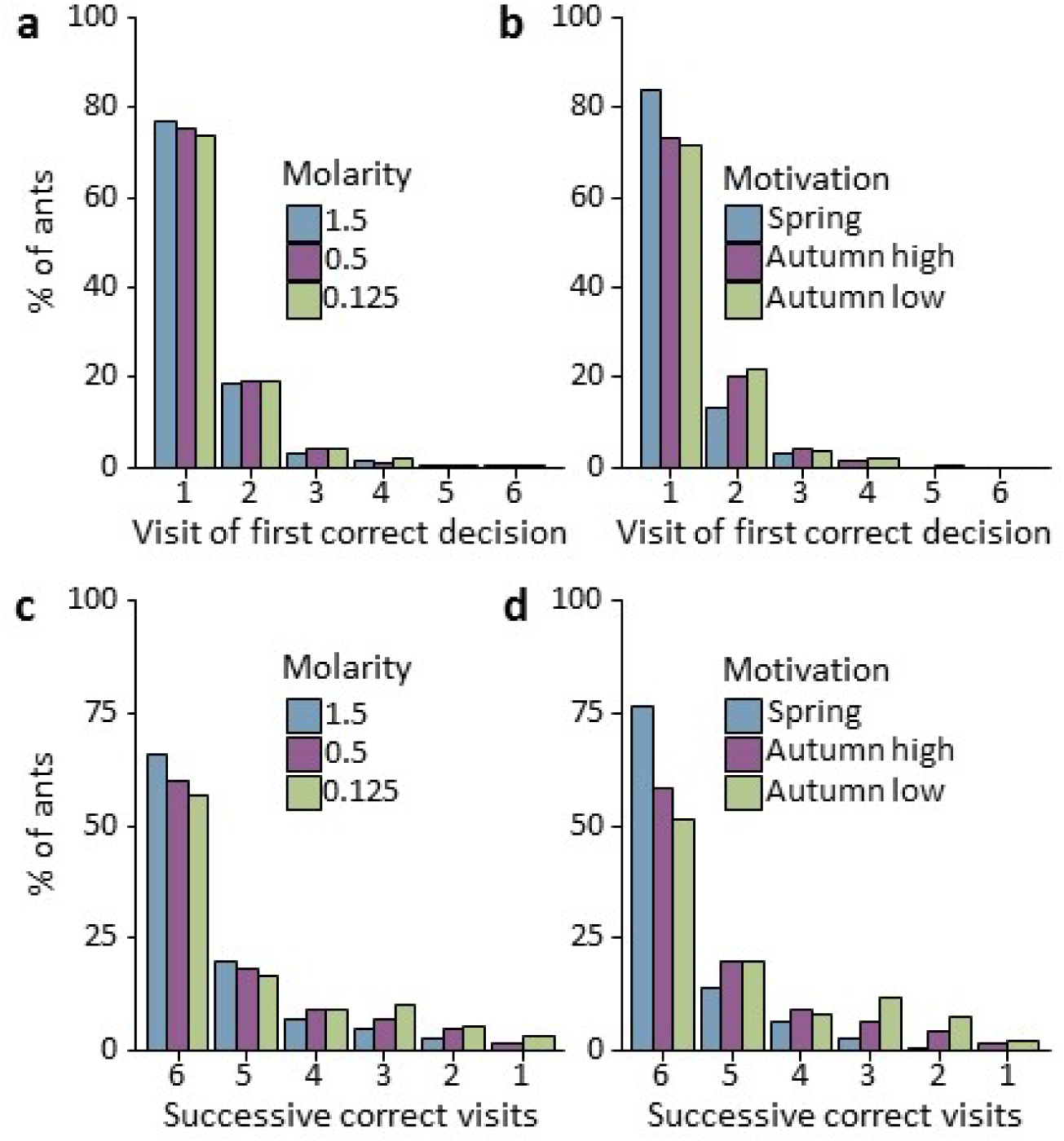
a, b: Percentage of ants making their first correct decision on the first down to the last visit. Most ants made their first correct decision on the first return visit to the food. While very small differences are seen between different food qualities (a), they become more prominent between different motivational levels (b). c, d: Percentage of ants making 6 correct visits in a row (100% correct) down to only 1 (no two correct visits in a row) as a measure of consistency of ant behaviour. While there is a trend for longer streaks with increasing food quality (c), motivational effects are stronger, with the majority of ants tested in spring making no error at all (d).Spring, Autumn high: 4 days of starvation; Autumn low: 1 day of starvation.

### Agent-based model results

The poor quality (0.125M) feeder was usually collectively avoided by the end of the model run (Fig. 4). When motivation levels were low or medium, the good quality (1.5M) feeder had the highest proportion of ants exploiting it. When motivation levels were high, however, colonies generally showed less “choosiness”, having a more equal distribution of foragers,with the majority of foragers exploiting the medium quality (0.5M) feeder. It should be noted that effect sizes are not large: at most 40% of ants exploited the most strongly chosen feeder, as compared to the null situation of one third.

**Fig. 4.**
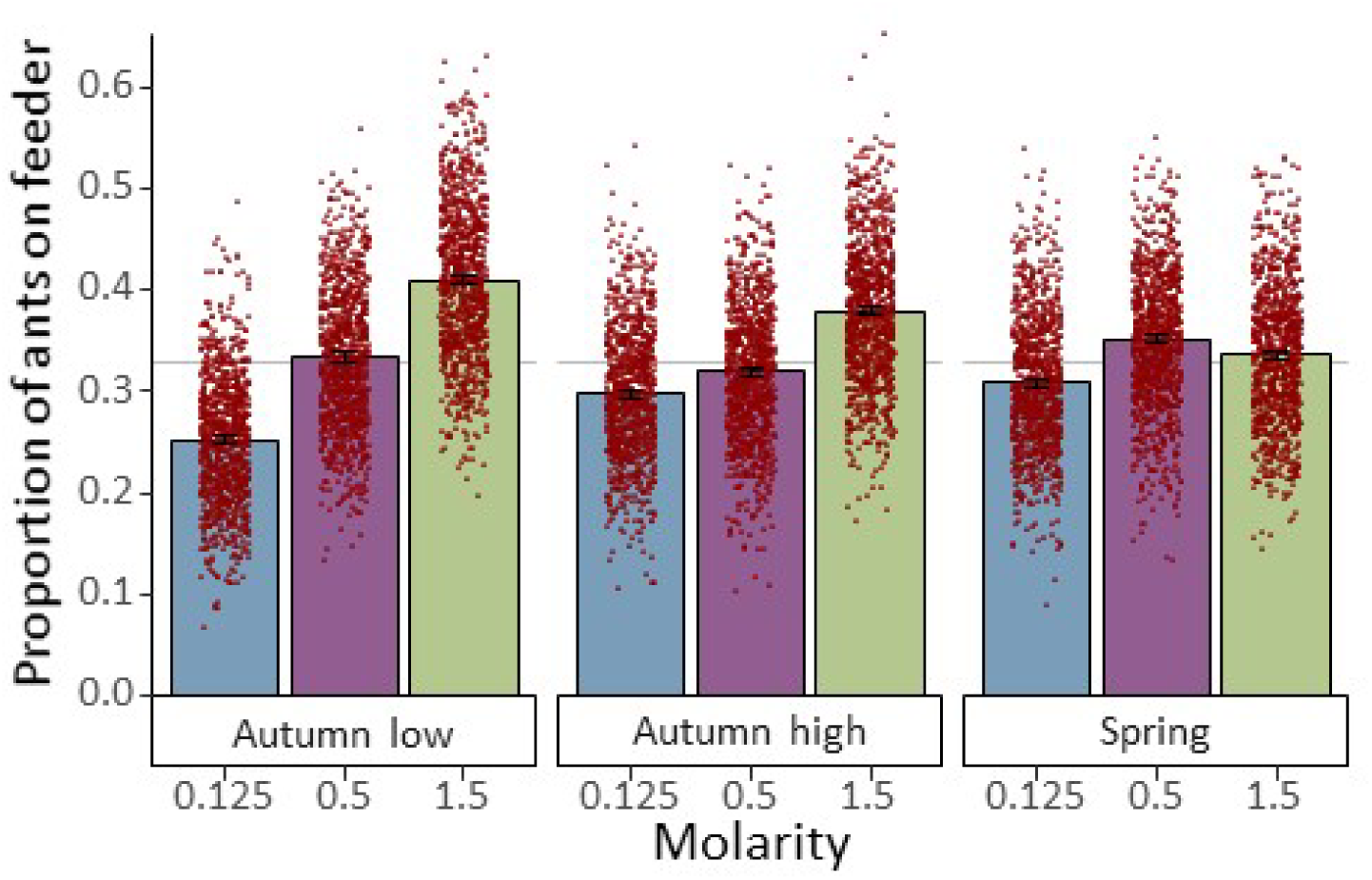
Proportion of agents exploiting each of the three feeders. These simulations used the “average ant” model of memory formation. Bars show means, error bars show 95% confidence intervals, dots show individual model run results. The dashed grey line shows the null situation of one third. For further model results, see OSM 2.

Increasing motivation level tended to increase the total food returned (due to more ants exploiting their memory rather than scouting), but the average quality of food returned to the nest was lower at higher motivation levels (see OSM 2 Fig. S2-2).

## Discussion

In our study, we investigated whether the effect of reward magnitude (sucrose concentration) and motivation (season and level of starvation) on differential learning is strong enough to explain apparent collective decisions of ants in the absence of communication.

In line with our hypothesis, the motivation of the ants significantly affected learning rates. Highest learning rates were found in spring, lowest in autumn in slightly starved colonies. However, surprisingly, the molarity of the reward did not significantly affect learning speed, although on average ants learned fastest when presented with the highest reward (Fig. 2a). Only a tendency towards higher sensitivity to food quality at lower motivation levels could be observed. Nonetheless, when these effects were built into an agent based model of ant foraging, colonies were found to send the largest proportion of foragers to the best food sources. This was especially the case at lower motivation levels: as foraging motivation drops, colonies become more collectively ‘choosy’. These findings suggest that at least part of an apparent collective decision to congregate at a high-quality food source could be accounted for by an as yet neglected annealing mechanism (Kirkpatrick et al. 1983; Černý 1985) based on learning: By preferentially memorising the best food source, ants will gradually improve until the optimum (all ants forage at the best food source) is reached, in the absence of communication.

It is noteworthy that even a weak and non-significant effect of food quality on learning could, in the agent based model, drive apparently ecologically sensible collective behaviour. We do not dispute that in laboratory experiments with mass-recruiting ants, collective feeder selection is driven mainly by pheromone deposition (Wilson 1962; Beckers et al. 1990, 1993), although memory undisputable can play a role in triggering collective behaviour (Czaczkes et al. 2016). However, such near-unanimous collective decisions do not well represent collective foraging on carbohydrate sources by ants in the wild (Devigne and Detrain 2005). In natural situations with limited, depleting but replenishing food sources, collective foraging is better described by models in which memory plays the biggest role in individual decision making (Czaczkes et al. 2015a). Understanding the effect of memory on foraging decisions is thus key to understanding real-world colony foraging decisions.

Overall, the learning rates of *Lasius niger* ants were very fast, with most ants (>70%) choosing the correct path already on the first visit after familiarization (Fig. 2a, b), resembling results of other studies (Grüter et al. 2011; Czaczkes et al. 2013). This highlights the ecological importance of route memory in *L. niger*, allowing the ants to find distributed, semi-permanent food sources repeatedly. In accordance with studies conducted on honey bees, decreasing rewards (1.5, 0.5, 0.125M) led to decreasing learning rates (e.g. Scheiner et al. 1999, 2004; 2005), although the differences were subtle and not significant in our study. Especially in highly starved ants, differences in learning rates were small, and these only became more prominent under low foraging motivation (Fig. S1). In our setup, however, it is impossible to distinguish between actual learning and persistence effects: A well-satiated ant might just alternate T-arms to explore. Moreover, more than 65% of the 1-day-starved ants did not perform all 6 runs when presented with the lowest, 0.125M, reward, but rather remained in the nest after some visits, as opposed to only ∼6% in highly starved autumn group and 0% in spring colonies (table 1). Under low starvation pressure, ants thus ignore poor quality food, which was also found in our agent-based model and is ecologically sensible (Seeley 1986). By this mechanism, at intermediate and low foraging motivation conditions, ant colonies will concentrate their foraging effort only on high quality food sources, with workers simply not returning to low quality feeders (table 1). This mechanism again relies on memory and persistence rather than differential recruitment.

Learning curves (Fig. 2a, b) are often used to present learning rates of groups, under the assumption that the group probability is a good representation of individual performance. This approach, however, neglects individual differences in learning rates (Pamir et al. 2011, 2014). Motivational differences between individuals are masked by overall performance and individuals seldom display a gradual improvement, but rather a binary response (Fig. 3 a-d, and Pamir et al. 2011, 2014). Furthermore, group level representations cannot show inconsistent behaviour of individuals, such as making errors after being correct before. We found such inconsistent behaviour in our study (Fig. 3c, d), with some ants in autumn, especially in the low motivation condition, not making two correct visits in a row. Nonetheless, the majority (75% in spring, >50% in autumn) of ants not only decided correctly on the first revisit, but also continued to be correct for the remaining 5 visits. These observations might be explained in terms of an exploration/exploitation trade-off (Cohen et al. 2007; Mehlhorn et al. 2015; Patrick et al. 2017). Animals have to decide between either exploiting available food or exploring the surroundings in order to find new food sources. In our experiment, starved ants readily exploited the available food and thus maximised their energy intake, while well satiated ants were more likely to explore (move to the other arm), especially when the encountered food was of low quality.

As the experimental setup only allowed individual ants to be tested, and completely preventing pheromone mediated recruitment in a whole colony is not technically feasible, we used an agent-based model (see OSM 1) to explore the impact of the observed learning rates on colony level. In the model, low motivation colonies mostly retrieved high quality food. This is sensible, as the processing and storage capacity of a colony is limited in such situations. By contrast, highly motivated colonies maximised food intake by collecting all food sources equally, leading to a decrease in energetic gain per load while maximising total energetic input. Such decreased choosiness when deprived of food can also be observed in other animals such as spiders (Pruitt et al. 2011). As the model was designed to explore the effects of differential learning on foraging, we purposefully did not include differential drop-out rates. Such a model would have to account for the costs of foraging, for which we lack good parameterisation. Including such an effect would most likely strengthen the patterns we see in our model: We would expect much greater ‘choosiness’, higher efficiency, and lower overall food intake at lower motivation levels.

The season of the year is known to affect foraging in social insects (Quinet et al. 1997; Ray and Ferneyhough 1997; Beekman and Ratnieks 2000; Scheiner et al. 2003; Mailleux et al. 2006; Cook et al. 2011, 2011; Al Toufailia et al. 2013), but effects on laboratory-reared colonies are less clear (Ray and Ferneyhough 1997). Our tested laboratory ant colonies clearly displayed decreased learning rates in autumn (Fig. 2b). In spring, *L. niger* is usually in dire need of food to begin worker production while being exposed to fluctuations in food supply, as the aphid colonies first need to establish (Mailleux et al. 2006). In such a variable environment, it is ecologically sensible to fully exploit all available sources. In autumn, ants usually stop egg production (Kipyatkov 1993) and activity and energy needs decrease with falling temperature (MacKay 1985). In our study, this variable response to food over the seasons seems to prevail in the laboratory colonies as well. Importantly, our queenless colonies are reinforced multiple times per year with workers from outside stem colonies, constituting a possible source of seasonal behaviour. Furthermore, we observe very large fluctuations in the amount of food consumed by colonies over the course of the year even in unreinforced colonies, with consumption rates falling dramatically towards autumn (FBO and TJC, pers. obs.).

To conclude, both reward quality and motivation can in principle lead to an adaptive increase in foraging efficiency via differential learning, without the need for any communication. This is in line with Dussutour and Nicolis (2013), who demonstrated that even ants which do not recruit to food sources can nonetheless focus on the best food source in a changing environment. They demonstrated, as we have, that such collective behaviour can be achieved by the increased retention of individuals to the best option, although their proposed retention mechanism relies on conspecifics and the quality of the food source while ours is based on memory and motivational effects. Both retention due to occupancy time at a food source and due to persistent visiting likely play a role in real-world situations. Such a collective decision without communication or comparison is conceptually identical to an annealing process and represents one of the simplest types of collective behaviour. Route memories alone have been demonstrated to be sufficient to trigger collective behavioural patterns in ants, such as becoming stuck in local foraging optima (Czaczkes et al. 2016). This study shows how memory could also play a role in the selection of the best resource. When studying collective decision-making, researchers tend to focus on interactive effects such as positive feedback and stigmergy. However, simpler non-interactive individual-level behavioural effects should also always be considered as an alternative, or contributing, mechanism driving group-level behaviour.

## Acknowledgments

We cordially thank Benedict Grüneberg for collecting part of the data and Florian Hartig for fruitful discussions about statistical methods.

## Ethical approval

All applicable international, national, and/or institutional guidelines for the care and use of animals were followed.

